# Lentiviral vectors transduce lung stem cells without disrupting plasticity

**DOI:** 10.1101/2020.10.19.345611

**Authors:** Ashley L. Cooney, Andrew L. Thurman, Paul B. McCray, Alejandro A. Pezzulo, Patrick L. Sinn

## Abstract

Life-long expression of a gene therapy agent likely requires targeting stem cells. Here we ask the question: does viral vector transduction or ectopic expression of a therapeutic transgene preclude airway stem cell function? We used a lentiviral vector containing a GFP or cystic fibrosis transmembrane conductance regulator (*CFTR*) transgene to transduce primary airway basal cells from human cystic fibrosis (CF) or non-CF lung donors and monitored expression and function after differentiation. Ussing chamber measurements confirmed CFTR-dependent chloride channel activity in CF donor cells. Immunostaining, quantitative real-time PCR, and single-cell sequencing analysis of cell-type markers indicated that vector transduction or CFTR expression does not alter the formation of pseudostratified, fully-differentiated epithelial cell cultures or cell type distribution. These results have important implications for use of gene addition or gene editing strategies as life-long curative approaches for lung genetic diseases.

## Introduction

Cystic fibrosis (CF) is caused by mutations in cystic fibrosis transmembrane conductance regulator (*CFTR*) which encodes an anion channel that contributes to regulation of airway surface liquid volume and composition. Without functional CFTR protein at the cell surface, dysregulated chloride and bicarbonate permeability ultimately leads to reduced innate immune defenses, bacterial colonization, inflammation, and mucus plugging that gradually and irreversibly destroy the lungs. Complementing or repairing *CFTR* in the appropriate pulmonary cell types early in life would prevent CF-related lung complications.

Many cell types in the conducting airways express *CFTR*, including: ciliated, non-ciliated, secretory, and ionocytes at the airway surface, as well as serous cells within the acini of the submucosal glands (1, 2). These cell types are obvious targets for CF gene therapy; however, they are often terminally differentiated and corrected CFTR expression would be lost as cells turnover. To achieve life-long correction from a gene therapy intervention, permanent genomic modification of self-renewing cells is likely required. Unlike many organs, the respiratory epithelium has multiple stem/progenitor cell populations. Different progenitor cell types are responsible for maintaining discrete niches from the trachea to the alveoli (3), and their roles may vary in health and disease. The heterogenous population of basal cells, which comprise the progenitor cells of the trachea and mainstem bronchi (4), have remarkable plasticity to respond to injury and repair the surrounding epithelia (reviewed in (5)).

Primary cultures of airway basal cells grown at an air-liquid interface mimic human airways and will reconstitute a pseudostratified epithelial sheet with multiple CFTR-expressing cell types (6, 7); however, basal cells do not typically express CFTR protein (8). The ramifications of exogenous expression of CFTR in basal cells are unknown and have important implications for gene therapy. Early evidence of CFTR expression in non-epithelial cells suggested altered metabolism, growth abnormalities, and potential consequences of exogenous transgene expression (9, 10). Since then, no significant progress has confirmed this and no studies have been performed using human primary basal cells. Here, we transduce a population of primary human basal cells with a lentiviral vector expressing either a GFP reporter gene or *CFTR* to address fundamental unanswered questions, including: 1) will lentiviral transduction or *CFTR* expression in basal cells alter their pluripotency and differentiated cell type distribution and 2) do CF cells complemented with *CFTR* have a global mRNA transcript expression profile that more closely resembles CF or non-CF patterns?

## Results

### Basal cell collection, transduction, differentiation, and phenotypic correction

Human lungs from multiple CF and non-CF donors were used for these studies. Primary basal cells were isolated from the trachea and bronchi as previously described (11) (**Figure 1A**). Cells isolated from both CF and non-CF airways expressed high levels of the known basal cell markers KRT5 (95%), p63 (92%), and NGFR (80%) and low levels of the ciliated cell marker, α-tubulin (3%) (**Figure 1B**). Basal cells were seeded onto Costar Transwells and transduced at the time of seeding with either HIV-GFP or HIV-CFTR (shown schematically, **Supplemental igure 1**) (MOI=5) overnight, left submerged for 2-3 days, then grown at an air-liquid interface (ALI) until well-differentiated (>4 weeks). Untransduced control cells were cultured in parallel. We quantified GFP in well-differentiated epithelial cells by flow cytometry to confirm that transgene expression was retained following differentiation. We observed 75-80% GFP positive cells in both CF and non-CF cultures (**Figure 1C**).

**Figure 1.**
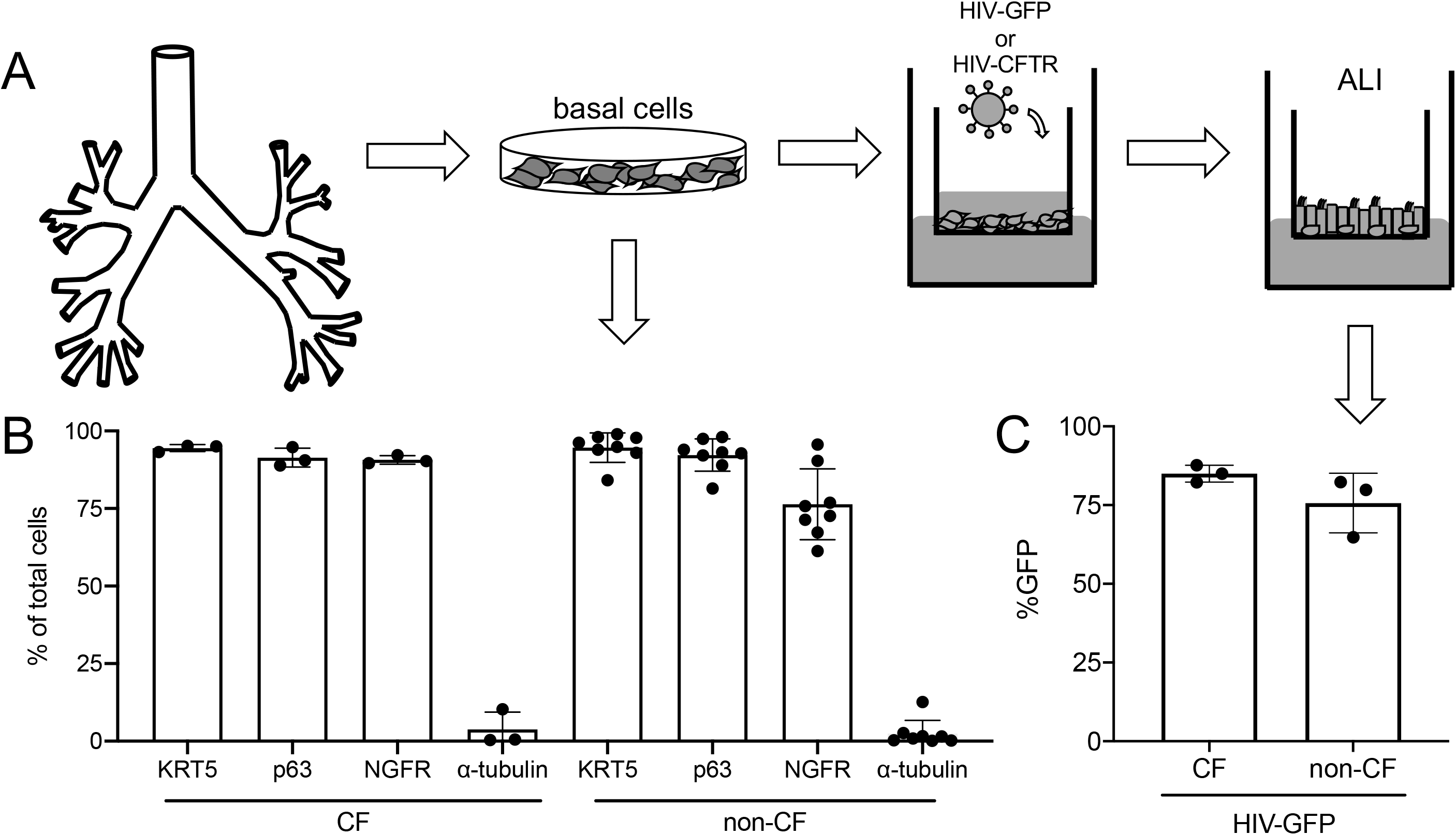
Primary basal cells express known markers and are readily transduced by a lentiviral vector. A) The experimental design model is shown. Primary basal cells are harvested from the trachea and mainstem bronchi and cultured until 80% confluent. Cells were then seeded on collagen-coated polycarbonate membranes and left untransduced or transduced overnight with HIV-GFP or HIV-CFTR. Cells were then grown submerged for 2-3 days when apical media is removed and cells are grown at an air liquid interface (ALI) until well-differentiated (>28 days). B) CF and non-CF Basal cells were assayed by flow cytometry for known basal cell markers KRT5, p63, and NGFR. α-tubulin was included as a negative control (CF n=3, non-CF n=8). C) GFP expression in well-differentiated epithelia was quantified by flow cytometry. (CF n=3, non-CF n=3).

To examine the morphology of cultured epithelia, immunostained cultures were imaged using confocal microscopy. Polarized airway epithelial cells had a pseudostratified columnar morphology with actin belts (phalloidin, white) and cilia (α-tubulin, red) (**Figure 2A-C**). XZ images (lower panels) revealed pseudostratification. As expected, GFP expression (green) was restricted to HIV-GFP treated cultures (**Figure 2B**). We next measured the anion transport properties of each condition. We hypothesized that complementing *CFTR* in CF basal cells using a lentiviral vector would achieve phenotypic correction after differentiation of transduced basal cells. As shown in **Figure 2D**, CF cells transduced with HIV-CFTR exhibited a greater change in short circuit current in response to the cAMP agonists forskolin and IBMX (F&I) and the CFTR inhibitor, GlyH-101, compared to untreated or HIV-GFP treated cells. These responses demonstrate complementation of the CF defect and were indistinguishable from non-CF cultures. Moreover, supplemental expression of *CFTR* by a lentiviral vector in non-CF cultures did not result in supraphysiologic changes in CFTR-dependent current.

**Figure 2.**
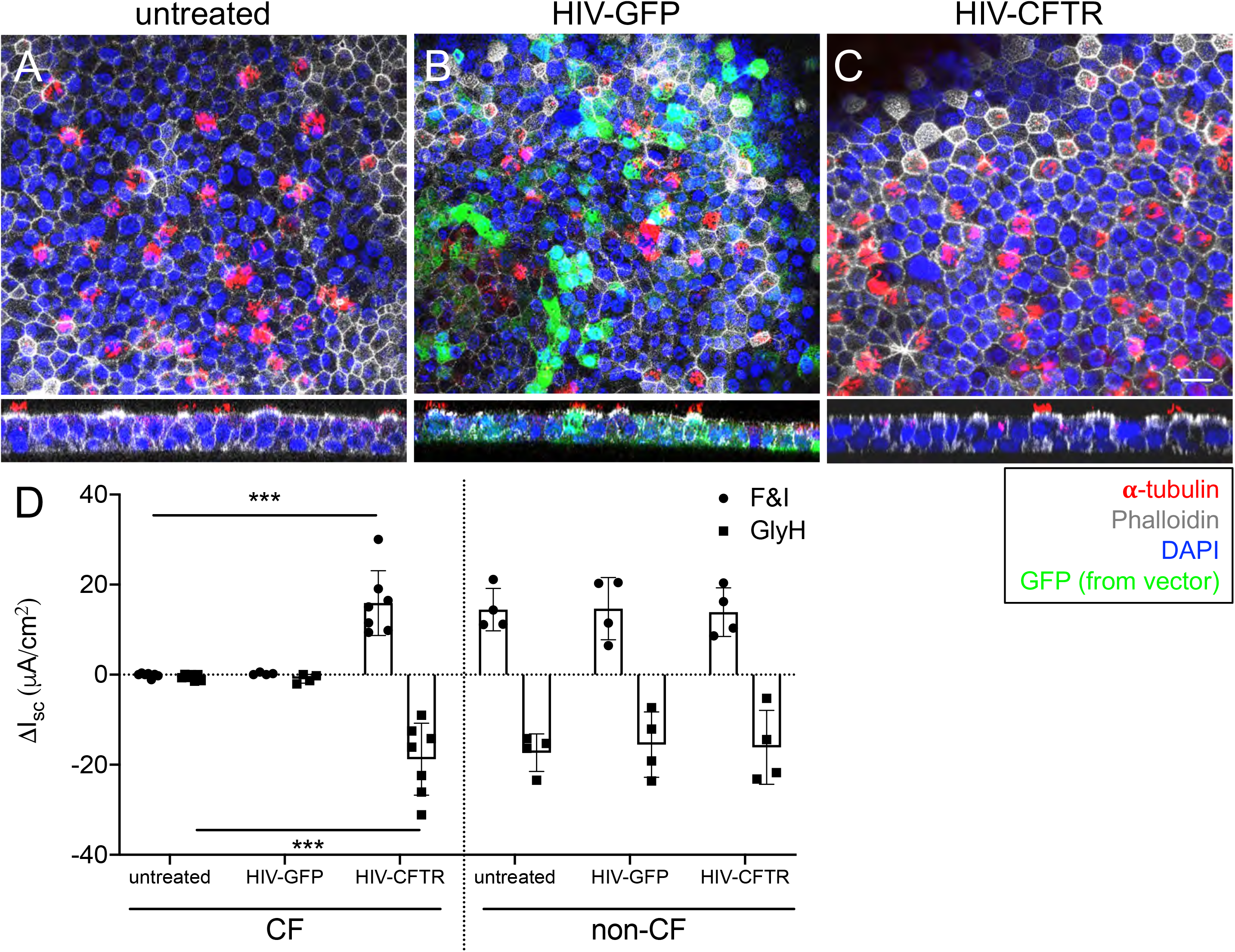
Primary basal cells differentiate into pseudostratified columnar epithelia and correct the CF anion channel defect. Well-differentiated cultures were immunostained for actin using Phalloidin (gray), cilia using α-tubulin (red), and nuclei by DAPI (blue). GFP expression is from the lentiviral vector. XY and XZ images are shown for each group. Representative images of are shown in panels A) untreated, B) HIV-GFP treated and C) HIV-CFTR treated. Scale bar = 10 μM. D) Ussing chamber analysis of airway epithelia. Data are presented as a change in short circuit current (ΔI_sc_) in response to F&I or GlyH. CF and non-CF cells were either left untreated or treated with HIV-GFP or HIV-CFTR (n=7 CF, n=4 non-CF).

### Single-cell transcriptome profiles of transduced primary basal cells following differentiation

To further analyze how basal cells respond to lentiviral transduction and CFTR expression, we used single-cell RNA (scRNA) sequencing to assess transcript levels by cell type. We compared multiple CF (n=4) and non-CF (n=3) donors. For each donor, we prepared a scRNA library and performed deep sequencing following 3 conditions (untreated, HIV-GFP, and HIV-CFTR). In total, 21 individual scRNA libraries were sequenced. We first confirmed that the appropriate cell types arose from basal cell differentiation, including secretory, basal, ciliated, ionocytes, and pulmonary neuroendocrine cells (PNECs) (2, 4, 12) **(Figure 3A**). Approximately 50% of the cells were secretory, 25% were basal, 25% were ciliated, and <1% were ionocytes or PNECs (**Figure 3B**). The major cell types (secretory, basal, and ciliated) appeared in similar ratios regardless of treatment (untreated, HIV-GFP, or HIV-CFTR) or disease (CF or non-CF). However, there were fewer ciliated cells in CF (~15%) than in non-CF (30%) cells. The 10 most highly expressed transcripts in each cell type are listed in a heat map (**Figure 3C**) where color intensity represents expression levels (yellow = high, black = low). The expression levels of canonical cell type markers were also visualized using a violin plot for secretory cells (BPIFA1), basal cells (KRT5), ciliated cells (FOXJ1), ionocytes (ASCL3), and PNECs (ASCL1) (**Figure 3D**).

**Figure 3.**
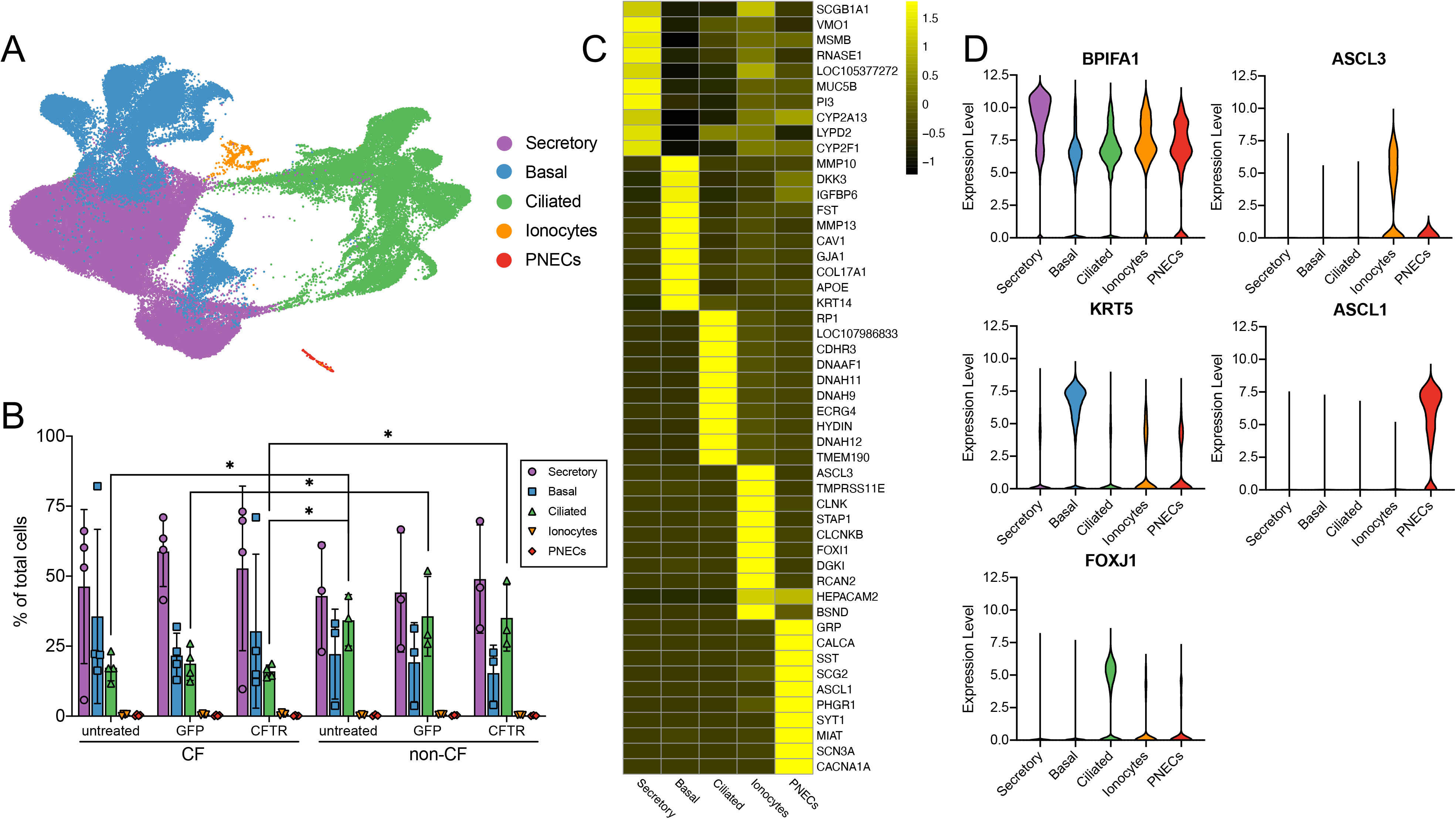
Summary of single cell RNA-seq. A) Single cell RNA sequencing revealed five cell types, shown via Uniform Manifold Approximation and Projection (UMAP). B) Cell type composition was quantified for each treatment in CF and non-CF cells with each point representing one donor. C) To validate labeling of cell types, scaled average expression of most highly upregulated genes are plotted by cell type. D) Distributions of normalized gene expression levels are shown for known cell markers BPIFA1 (secretory cells), ASCL3 (ionocytes), KRT5 (basal cells), ASCL1 (PNECs), and FOXJ1 (ciliated cells) (n=4 CF, n=3 non-CF).

As further confirmation of appropriate expression levels of cell type markers, we used quantitative real-time PCR (qRT-PCR) to determine the relative abundance of *p63* (basal cells), *FOXJ1* (ciliated cells), *MUC5AC* and *MUC5B* (secretory cells) markers compared to undifferentiated basal cells in the cultures (**Supplemental Figure 2A-D**). Relative to the initial expression in basal cells, we observed similar levels of *Pan*-Δ*Np63*, a marker for basal cells, and high levels of *FOXJ1*, *MUC5AC*, and *MUC5B*, confirming well-differentiated airway epithelia (13). Consistent with previous reports, CF cells expressed higher levels of *MUC5AC* and *MUC5B* transcripts than non-CF cells, possibly due to goblet cell metaplasia (14). These levels were unchanged when CFTR was expressed in CF cells. These data suggest that the expression profile of non-CF and CF airway epithelial basal cells may vary. Moreover, the minimal change in pattern observed with exogenous *CFTR* expression suggests the genotype-associated difference may not be determined by CFTR function.

Once we observed that basal cells from all groups differentiated into the appropriate cell types and ratios, we next asked if the expression profile of CF cells treated with HIV-CFTR more closely resembled CF or non-CF cells. Principal Component Analysis (PCA) plots demonstrated that CF + HIV-CFTR (stars) clustered more closely with CF (squares) than non-CF (circles) (**Figure 4**). This is somewhat expected since they are paired from the same donor; however, CF + HIV-CFTR from all 4 donors cluster more closely with non-paired CF donors than non-CF cells.

**Figure 4.**
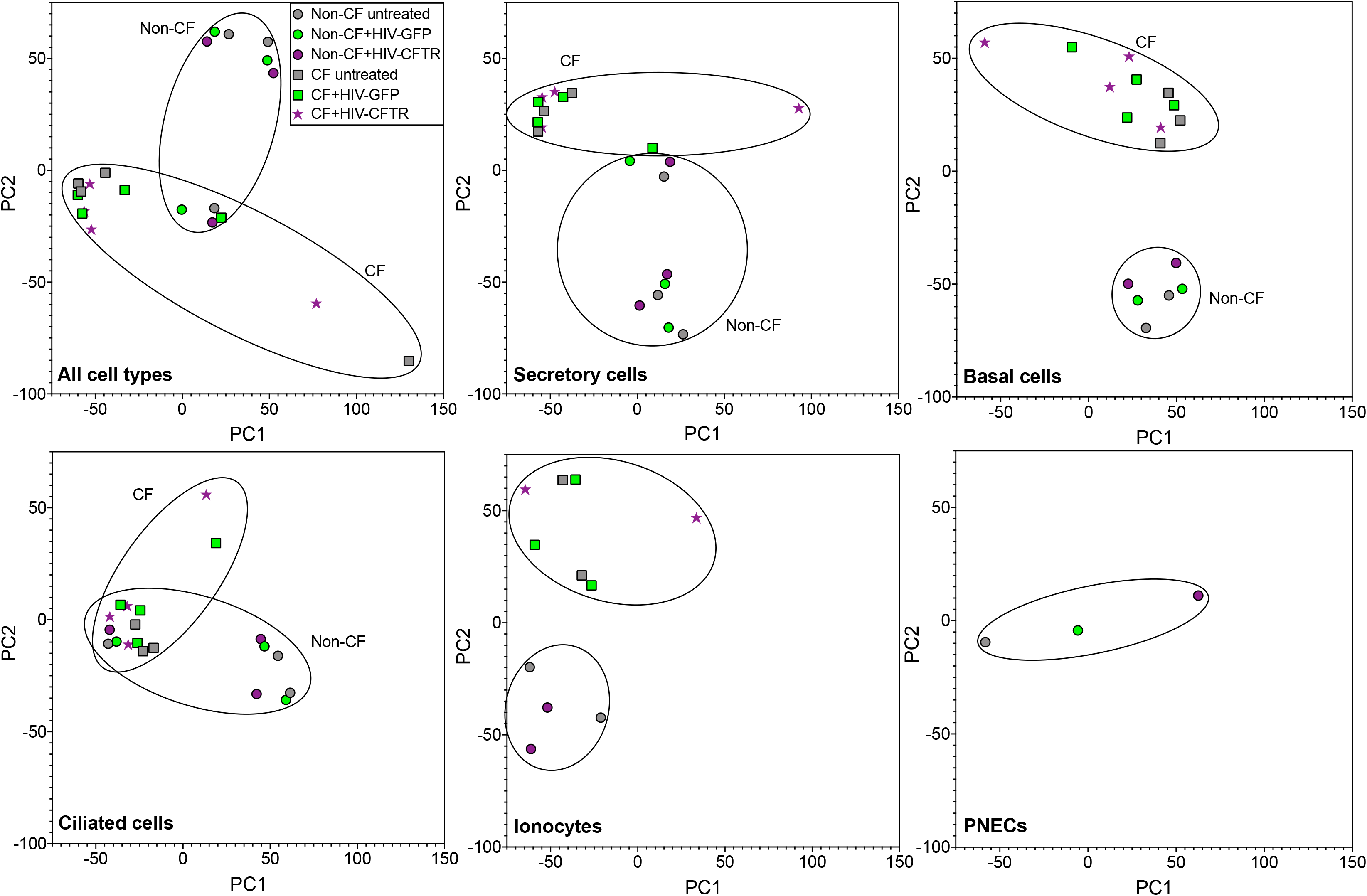
Transcriptomic similarity of cultures. For single cell RNA-seq data, cells were stratified by cell type, gene expression profiles were summarized for each culture, and principal component scores were computed. Plots of the first two principal components (PC1 and PC2) show gene expression similarity between cultures by disease and treatment (n=4 CF, n=3 non-CF).

Changes to the transcriptome following lentiviral vector integration or exogenous *CFTR* expression could reveal the impact of gene therapy on cellular gene expression. Here, we analyzed genes differentially expressed in both CF and non-CF cells following treatment with a lentiviral vector expressing either GFP or CFTR relative to their untreated controls. We observed few differentially expressed genes. In each case, we analyzed differentially expressed transcripts among all cell types combined (**Figure 5**) as well as individual cell types (**Supplemental Figure 3**) as shown by volcano plots. The X-axis indicates the relative expression levels between each group (GFP vs. untreated or CFTR vs. GFP) and the Y-axis shows the P value significance. The majority of the differentially expressed genes originated from the lentiviral cassette in transduced cells (i.e. GFP or the Ankyrin insulator element) (black dots, **Figure 5A, C**). In CF cells treated with HIV-CFTR, a small number of genes were slightly above the threshold of *FDR* > 0.05 (**Figure 5B**). Of note, non-CF cells treated with HIV-CFTR had no significantly differentially expressed genes when compared to HIV-GFP treated cells (**Figure 5D**). Examples of non-vector related genes include *CCL2, IL6, PRSS2,* and *PRSS37. CCL2* and *IL6* are a chemokine and cytokine, respectively; *PRSS2* and *PRSS37* belong to the serine protease family. A complete list of all differentially expressed endogenous genes is available in **Supplemental Table 1**. The low number of differentially expressed transcripts suggests that delivery of an exogenous transgene by a lentiviral vector has minimal effects on the host transcriptome. Moreover, they suggest that *CFTR* expression and activity regulates very few genes transcriptionally.

**Figure 5.**
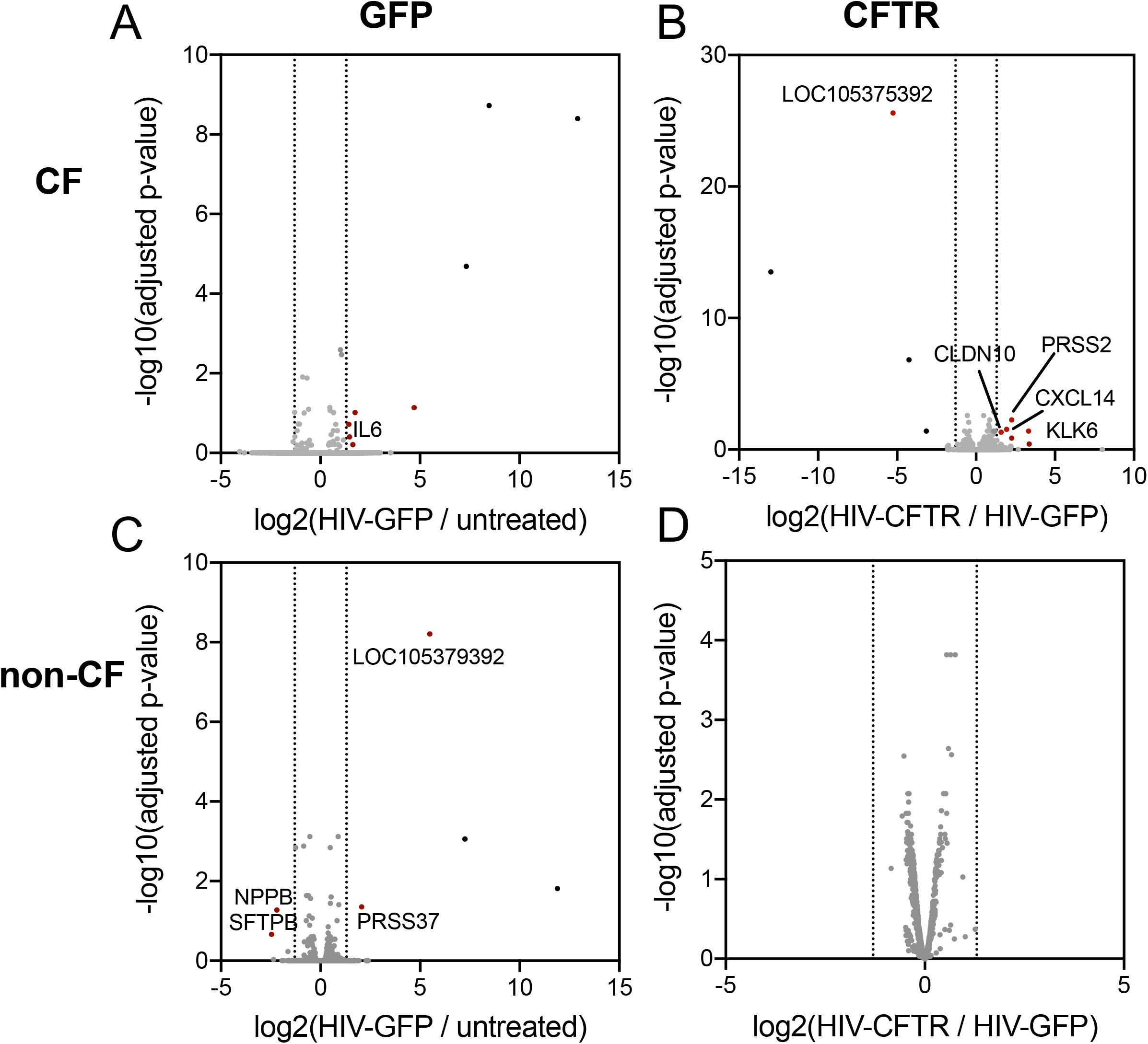
*Effects of transduction and* CFTR *correction on gene expression*. Differential expression analysis was performed to visualize the effect of HIV-GFP transduction (A) CF, C) non-CF), and HIV-CFTR transduction (B) CF, D) non-CF) on gene expression for each disease group. Volcano plots show fold change in expression (log2 fold change) versus statistical significance (−log10(pval)). HIV-GFP (positive log2 fold change) gene expression was compared to untreated cells (negative log2 fold change). Red dots indicate differentially expressed genes, gray dots indicate non-differentially expressed genes, and black dots indicate vector encoded genes. For CFTR complementation, gene expression was compared between HIV-CFTR (positive log2 fold change) and HIV-GFP (negative log2 fold change). Black dots indicate vector encoded genes. Notable genes are labeled (n=4 CF, n=3 non-CF).

We next confirmed that the delivered transgenes were expressed in all cell types As expected, we observed GFP expression in the HIV-GFP transduced cells by qRT-PCR (**Figure 6A**). An example of the scRNA sequencing alignment coordinates shows the number of reads at each nucleotide position within GFP (**Figure 6B**). This indicates that the GFP reads detected were present in GFP samples only. We observed GFP expression in all cell types including secretory, basal, ciliated, ionocytes, and PNECs by scRNA seq (**Figure 6E**). These data indicate that all cell types appropriately differentiated from a population of basal cells transduced with a lentiviral vector expressing GFP and that using a lentiviral vector to transduce airway stem cells does not preclude stem cell differentiation.

**Figure 6.**
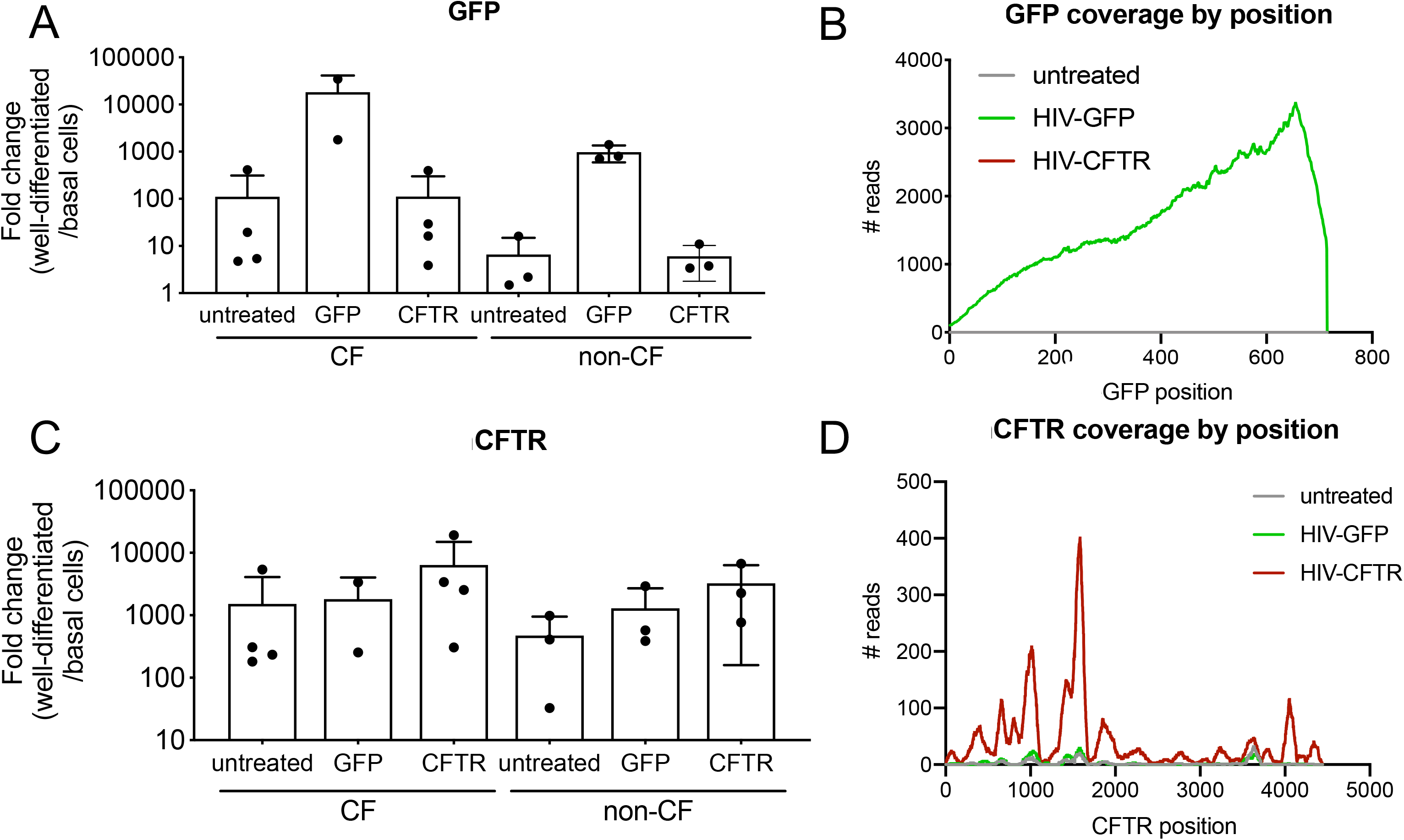
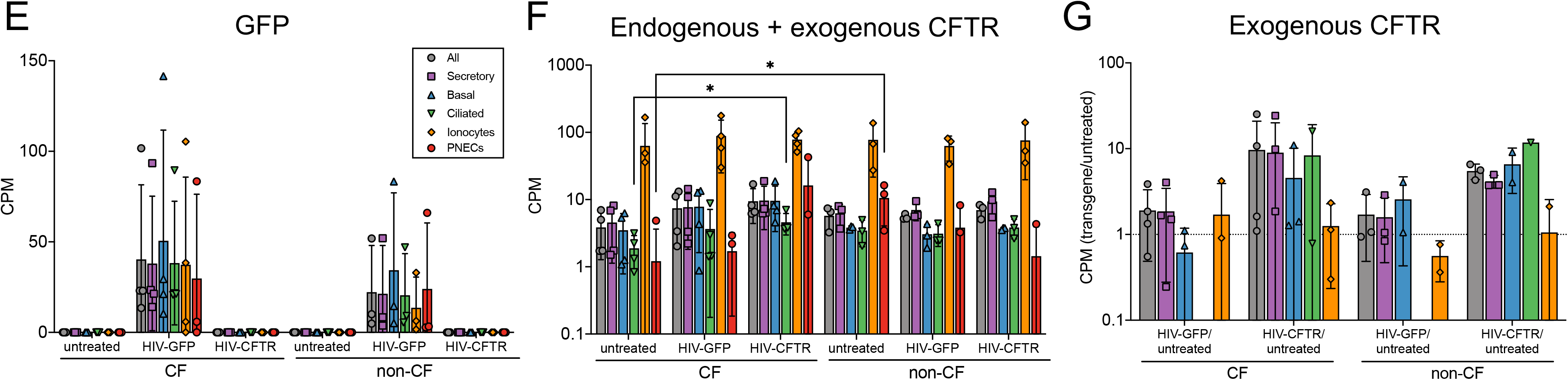
*GFP and* CFTR *expression levels in airway epithelia by qRT-PCR and single-cell sequencing*. A) Real Time quantitative PCR levels of GFP is shown for the indicated treatment groups. Fold-change is expressed as well-differentiated over basal cells for each marker (n=5 CF, 4 non-CF). B) For one CF culture, single cell RNA-seq coverage of GFP transgene (eGFP). Coverage is shown for untreated, HIV-GFP, and HIV-CFTR treatments. C) Real Time quantitative PCR levels of *CFTR* is shown for the indicated treatment groups. Fold-change is expressed as well-differentiated over basal cells for each marker (n=5 CF, 4 non-CF). D) For one CF culture, single cell RNA-seq coverage of *CFTR* transgene (CFTR). Coverage is shown for untreated, HIV-GFP, and HIV-CFTR treatment groups (n=4 CF, n=3 non-CF). E) Gene expression levels are shown in counts per million (CPM) of GFP for each human donor and treatment group. F) Gene expression levels of total CFTR (endogenous + CFTR transgene) for each donor and treatment group are shown. G) Relative levels of exogenous CFTR transgene expression is presented as HIV-GFP over untreated and HIV-CFTR over untreated. Dotted line at Y=1 indicates expression levels equivalent to basal cells. Legend indicating cell types is shared among E, F and G (n=4 CF, n=3 non-CF).

We measured total *CFTR* mRNA expression by qRT-PCR and observed a trend towards higher levels in the HIV-CFTR treated cells than the HIV-GFP or untreated conditions (**Figure 6C**). We next sought to differentiate between endogenous *CFTR* mRNA and transcripts supplied by the vector. Representative alignment coordinates show the number of reads that match the input sequence (**Figure 6D**). We observed an increase in total (endogenous and *CFTR* transgene) *CFTR* transcripts in CF and non-CF cells treated with HIV-CFTR (**Figure 6F**). We next compared *CFTR* transgene levels from GFP or CFTR treated cells relative to untreated CF cells and observed that *CFTR* transcripts were increased in secretory, basal, and ciliated cells in HIV-CFTR transduced cells only (**Figure 6G**). In CF cells treated with HIV-CFTR, *CFTR* transcript levels were restored to non-CF levels in ciliated cells. This is consistent with the differential gene expression data in **Supplemental Figure 3 (CF, Ciliated cells)**. Also, PNECs seemed to show less *CFTR* in untreated CF cells than untreated non-CF cells. Interestingly, ionocytes expressed the highest *CFTR* levels but expression was not increased following the delivery of HIV-CFTR (**Figure 6G**). In summary, modest detectable increases in *CFTR* transcripts were sufficient to restore functional correction to CF epithelia. Additionally, *CFTR* expressed in basal cells is retained in basal cells and expressed in all major cell types after differentiation.

## Discussion

For a gene therapy treatment to correct CF airway disease for the life of a person, stable expression of a functional CFTR protein in surface airway epithelial cells is likely necessary. Of the many proposed gene therapy approaches that may satisfy this benchmark, lentiviral delivery of a constitutively expressed *CFTR* cDNA to airway progenitor cells is among the products with near term translational potential. Basal cells are the progenitor cells of the large airways; however, prior to this study, important questions about the consequences of *CFTR* expression in basal cells were unaddressed. Although mouse studies suggest that lentiviral vectors can persistently express CFTR for the life of the animal (15, 16), evaluating a gene therapy treatment specifically in human airway progenitor cells is crucial. Here, we analyzed primary basal cells from CF and non-CF donors and their responses to lentiviral vectors expressing GFP or CFTR. Our results suggest that neither a lentiviral vector nor *CFTR* expression significantly altered the differentiation potential of primary basal cells acquired from human donors.

Little is known about how restoring *CFTR* in airway epithelia affects the transcriptome. Small molecule correctors and potentiators, such as Ivacaftor and Trikafta, are beginning to inform us of the long-term effects of restoring CFTR activity (17, 18). Transcriptome profiling, including single-cell RNA sequencing, is a robust approach that can address fundamental questions about cellular responses to *CFTR* complementation and provide clues to underappreciated CF defects. Here we evaluated well-differentiated airway epithelia derived from primary human basal cells using single-cell sequencing. We found that lentiviral vector transduction of basal cells was associated with very few differentially expressed genes following differentiation. Supplemental *CFTR* expression in either CF or non-CF cells resulted in only minor transcriptomic changes, regardless of cell type.

A potential hurdle for either gene addition or gene editing strategies is access to lung progenitor cells. Basal cells line the basement membrane of the conducting airways and repopulate the surface epithelium following cell turnover or injury (4). Topical aerosol delivery deposits gene therapy vectors on the luminal surface of the airways, where access to basal cells is limited by tight junctions. However, we and others have shown that agents such as the natural airway surfactant lysophosphatidylcholine (LPC) (19–21) and calcium chelator EGTA (22) transiently increased epithelial permeability and allow access to the basolateral surface of columnar epithelial cells as well as basal cells. Additionally, some basal cell extensions may reach the luminal surface (23).

The percentage of basal cells is highest in the trachea (about 30%) and gradually decreases throughout the airway tree, reaching less than 6% in the proximal small airways (24). Progenitor cells of the small airways lie at the bifurcation of the bronchiolar and respiratory epithelium (25), and include club cells and alveolar type II cells, respectively. A major difference between stem/progenitor cells in the large versus small airway is accessibility from the lumen. The small airways are an important target for CF gene therapy and progenitor cells of this region are accessible without the need to disrupt tight junctions. We have previously shown that an aerosol delivery of vectors transduces both proximal and distal airways of a large animal model (21).

Delivering *CFTR* by a lentiviral vector to CF cells complements the anion channel defect (15, 16, 26, 27). Supraphysiologic current changes were previously observed with *CFTR* delivered to fully-differentiated primary cultures using an adenoviral-based vector. In that setting, mislocalization of CFTR to the basement membrane resulted in a net loss of anion transport (28). In our studies, Supplemental *CFTR* expression by a lentiviral vector showed no evidence of aberrant expression based on bioelectric properties. Transcriptome analysis revealed no consequential differences between untreated, GFP-treated, or CFTR-treated non-CF cells.

One observation consistent among all conditions was the ratio of cell types that arose from basal cells. These data suggest an intrinsic ability for progenitor cells to maintain an appropriate proportion of cell types that comprise the pseudostratified epithelium. Representing one of the smallest populations of cells in our dataset, ionocytes expressed the highest levels of *CFTR* expression, consistent with previous reports (2, 29). Importantly, while expression of *CFTR* in non-CF secretory, basal, and ciliated cells increased mRNA transcript levels, this did not result in an increase in CFTR-dependent Cl^−^ current. This could suggest a mechanism to regulate *CFTR* levels among various cell types. Secretory, basal, and ciliated cells from *CFTR*-transduced cells expressing *CFTR* indicate that basal cells can indeed differentiate into other cell types while retaining *CFTR* expression, further suggesting that *CFTR* expression does not disrupt stem cell plasticity.

We hypothesized that restoring *CFTR* would modify the transcriptome to mirror a non-CF cell-like state, but instead found that transcripts from CF cells complemented with HIV-CFTR more closely resembled CF cells than non-CF. These findings were consistent in the scRNA sequencing data as well as qRT-PCR. Through these studies, we compiled a list of non-vector encoded genes that were upregulated or downregulated in response to restoring *CFTR*. Understanding the impact that the expression of these genes have will require further investigation. Another interesting observation was that restoring *CFTR* to CF or non-CF cells increased transcript levels of *CFTR* in all cell types except ionocytes. We postulate that there may be an upper physiological limit of expressing CFTR in ionocytes, but this has yet to be investigated.

A primary objective of this study was to quantify the effects of *CFTR* expression on the pluripotency of airway progenitor basal cells. We transduced pure populations of human basal cells collected from multiple CF and non-CF human donors. This is a relevant and representative *in vitro* model system; however, this study has limitations. Because this is an *in vitro* model of epithelial cells derived from the trachea and mainstem bronchi, we did not evaluate immune responses to the vector or vector components, basal cell transduction efficiencies in an animal model, or progenitor cell populations of the submucosal glands, small airways, or alveoli. We did not measure persistence through basal cell proliferation for the following reasons: 1) passaging basal cells leads to senescence after 6-8 passages (13, 30), 2) cell turnover *in vitro* is not representative of an *in vivo* setting (31), and 3) immune responses *in vivo* may play a role in selective cell-mediated clearance.

Here we addressed the differentiation potential of *CFTR*-expressing basal cells compared to untreated or GFP-expressing cells in CF and non-CF primary cells. Based on qRT-PCR, morphology, physiologic assays, and single-cell RNA sequencing analyses, we provide evidence that lentiviral-delivery of *CFTR* to basal cells does not preclude formation of a well-differentiated airway epithelium with complementation of CFTR anion channel activity.

## Material and Methods

### Ethics statement

Basal cells from human CF and non-CF donors were isolated from discarded tissue, autopsy, or surgical specimens. Cells were provided by The University of Iowa *In Vitro* Models and Cell Culture Core Repository. We were not provided with any information that could be used to identify a subject. All studies involving human subjects received University of Iowa Institutional Review Board approval.

### Viral vector production

VSV-G pseudotyped HIV viral vectors with a PGK promoter driving either GFP (cloned by Laura Marquez Loza) or CFTR expression were produced at the University of Iowa Viral Vector Core (http://medicine.uiowa.edu/vectorcore/) using a four plasmid transfection method as previously described (32). The lentiviral vector was previously described and provided by Stefano Rivella (33). Lentiviral vectors were titered by droplet digital PCR (34), and/or by flow cytometry as part of the Vector Core service.

### Cell culture

Basal cells were cultured in LifeLine BronchiaLife (LifeLine Cell Technology, Carlsbad, CA) media for 3-5 days following lung harvest. When the cells reached 80% confluency they were washed in Phosphate Buffered Saline (PBS), lifted in TrypLE (Gibco, Gaithersburg, MD), and counted. 1×10^5^ cells were seeded on each 0.33 cm^2^ collagen IV coated polycarbonate transwell inserts (Corning Costar, Cambridge, MA) in Ultraser G (USG). Cultures were left untransduced or transduced with either VSVG-HIV-PGK-GFP or VSVG-HIV-PGK-CFTR overnight at a MOI of 5 at the time of seeding. Transwells remained submerged in USG for 2-3 days post-seeding, apical media was removed, and cells were grown at an air-liquid interface (ALI) until well-differentiated (>28 days).

### Flow cytometry

Following differentiation of the cultures, GFP was quantified by flow cytometry. Briefly, cells were stained using a LIVE/DEAD Fixable Stain (ThermoFisher Scientific, Waltham, MA), lifted in Accutase at 37°C for 15 minutes, and run through an Attune NxT Flow Cytometer (ThermoFisher Scientific, Waltham, MA) and percentage of GFP positive cells were calculated. Basal cells were assayed for basal cell markers Krt5 (ab193894; 1:600, Abcam, Cambridge, MA), p63 (ab246728; 1:600, Abcam Cambridge, MA), NGFR (345110; 1:600, BioLegend, San Diego, CA), and α-tubulin (NB100-69AF405, 1:300, Novus, Centennial, CO). Cells were treated with Foxp3/Transcription Factor Staining Buffer Set (ThermoFisher Scientific, Waltham, MA) according to manufacturer’s recommendations and stained for 1 hour at 4°C using the antibodies listed above. Cells were run on the Attune NxT Flow Cytometer and expression was gated on live cells.

### RNA isolation and qRT-PCR

Basal cells or well-differentiated epithelia were treated with Trizol and RNA was isolated using the Zymo Directzol Miniprep isolation kit (Zymo Research, Irvine, CA) according to the manufacturer’s instructions. cDNA was generated using Applied Biosystems High-Capacity cDNA Reverse Transcription Kit (ThermoFisher Scientific, Waltham, MA). qRT-PCR was performed with Power SYBR Green Master Mix (ThermoFisher Scientific, Waltham, MA). Primer sets are listed in **Supplemental Table 2**. Fold change was calculated as change in gene expression of well-differentiated epithelia over untreated basal cells.

### Electrophysiology

Short circuit current was measured in well-differentiated airway epithelia derived from primary basal cells from CF and non-CF donors. Conditions included untreated, HIV-GFP, or HIV-CFTR treated cells. Airway cultures were mounted in the Ussing chamber and their bioelectric properties were quantified as previously reported (26). Cells were pre-treated with forskolin and 3-Isobutyl-1-methylxanthine (IBMX) (F&I) overnight prior to Ussing chamber analysis. Assay protocol is as follows: Amiloride, 4,4’-Diisothiocyano-2,2’stilbenedisulfonic acid (DIDS), F&I, and GlyH-101 (GlyH). Results are reported as a change in short circuit current (ΔI_sc_) in response to F&I or GlyH.

### Single-cell RNA sequencing

Libraries were generated for single-cell RNA sequencing according to the Chromium Single Cell Gene Expression v3 kit 10X Genomics protocol (10X Genomics, Pleasanton, CA). ~5000 cells were combined with Gel Beads, Master Mix, and Partitioning Oil and loaded into a Chromium Next GEM Chip. Single cells were partitioned in oil to generate gel beads in emulsion (GEMs). GEMs were then dissolved and barcoded with Illumina TruSeq sequencing primer, barcode, and unique molecular identifier (UMI). A reverse transcription step next generated full-length cDNA. After an additional round of cDNA amplification, cDNA underwent enzymatic fragmentation to select the appropriate amplicon size and then labeled via End Repair, A-tailing, Adaptor Ligation, and PCR to construct final single-cell libraries for sequencing. Sequencing was performed on the HiSeq or NovaSeq 6000 platform.

### Bioinformatic Analyses

Raw sequencing reads were processed using CellRanger version 3.0.2 with alignment to a hybrid genome consisting of human genome reference GRCh38.p13 and the GFP and *CFTR* transgenes. Gene-by-cell count matrices were processed with the R package Seurat version 3.1.1 (35). Counts for each cell were normalized by total UMIs and log transformed to quantify gene expression for each cell. To reduce the data dimensionality for clustering and visualization, centered and scaled gene expression for the 2,000 mostly highly variable genes were further reduced to the first 20 principal component scores for input to a shared nearest neighbor clustering algorithm. Cell type identities were associated with each cluster by identifying upregulated genes in each cluster using a Wilcoxon rank sum test and comparing upregulated genes to a list of known airway epithelial markers.

### Deposited data

Single cell RNA sequencing data has been deposited in the GEO with accession number GSE159056.

### Statistics

Statistical comparisons were performed with cultures as the units of analysis. For each culture, cells were stratified by cell type, and gene counts were summed across cells, resulting in a gene-by-culture count matrix for each cell type. Comparisons of different cell proportions between CF and non-CF cultures and comparisons of gene expression levels for individual genes were performed using a t-test. Genome-wide differential gene expression analyses for the effect of VSVG-HIV-PGK-GFP and VSVG-HIV-PGK-CFTR transductions were performed using the R package DESeq2 version 1.22.2 (36) (R Core Team (2020). R: A language and environment for statistical computing. R Foundation for Statistical Computing, Vienna, Austria, https://www.R-project.org/). Differentially expressed genes were defined as those having an *FDR* < 0.05 and at least a two-fold difference in expression.

## Acknowledgements

We thank Bo Ram Kim, Guillermo Romano Ibarra, Laura Marquez Loza, Ian Thornell, Chris Wohlford-Lenane, and Brajesh Singh for technical assistance. We thank Phil Karp and Ping Tan in the In Vitro Models and Cell Culture Core for providing the basal cells from the many lung donors used in these studies. We thank Laura Marquez Loza, Amber Vu, Christian Brommel, Cami Hippee and Miguel Ortiz for their critical review of this manuscript. This work was supported by the NIH (P01 HL51670, P01 HL091842, P01 HL152960, R01 HL133089), the Cystic Fibrosis Foundation (COONEY18F0, SINN19XX0), The University of Iowa Center for Gene Therapy (DK54759), and the Roy J. Carver Chair in Pulmonary Research (P.B.M.).

## Author Contributions

ALC: designing experiments, data collection, analyzing and interpreting results, writing the manuscript

AT: data collection, analyzing and interpreting results, writing the manuscript

PBM: designing experiments, analyzing and interpreting results

AP: designing experiments, analyzing and interpreting results

PLS: designing experiments, analyzing and interpreting results, writing the manuscript

---

**Supplemental Figure 1.**
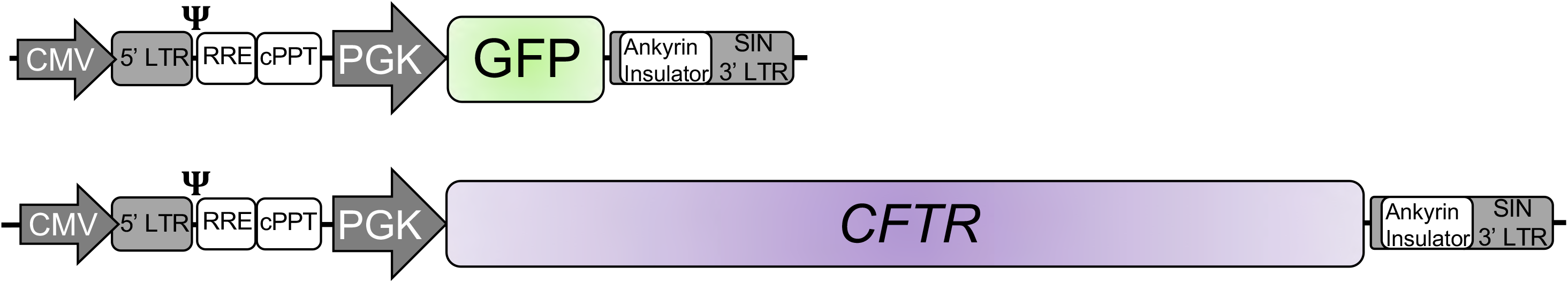
Schematics of integrated form of lentiviral vectors. Self-inactivating (SIN) lentiviral vector transgene cassettes include a CMV promoter driving the partial 5’ LTR (R, U5 region), packaging signal (Ψ), Rev-response element (RRE), and central polypurine tract (cPPT). The phosphoglycerate kinase (PGK) promoter drives either GFP or CFTR expression followed by the partial U3 region, Ankyrin insulator element, SIN 3’ LTR (R, U5 region). Elements shown are not to scale.

**Supplemental Figure 2.**
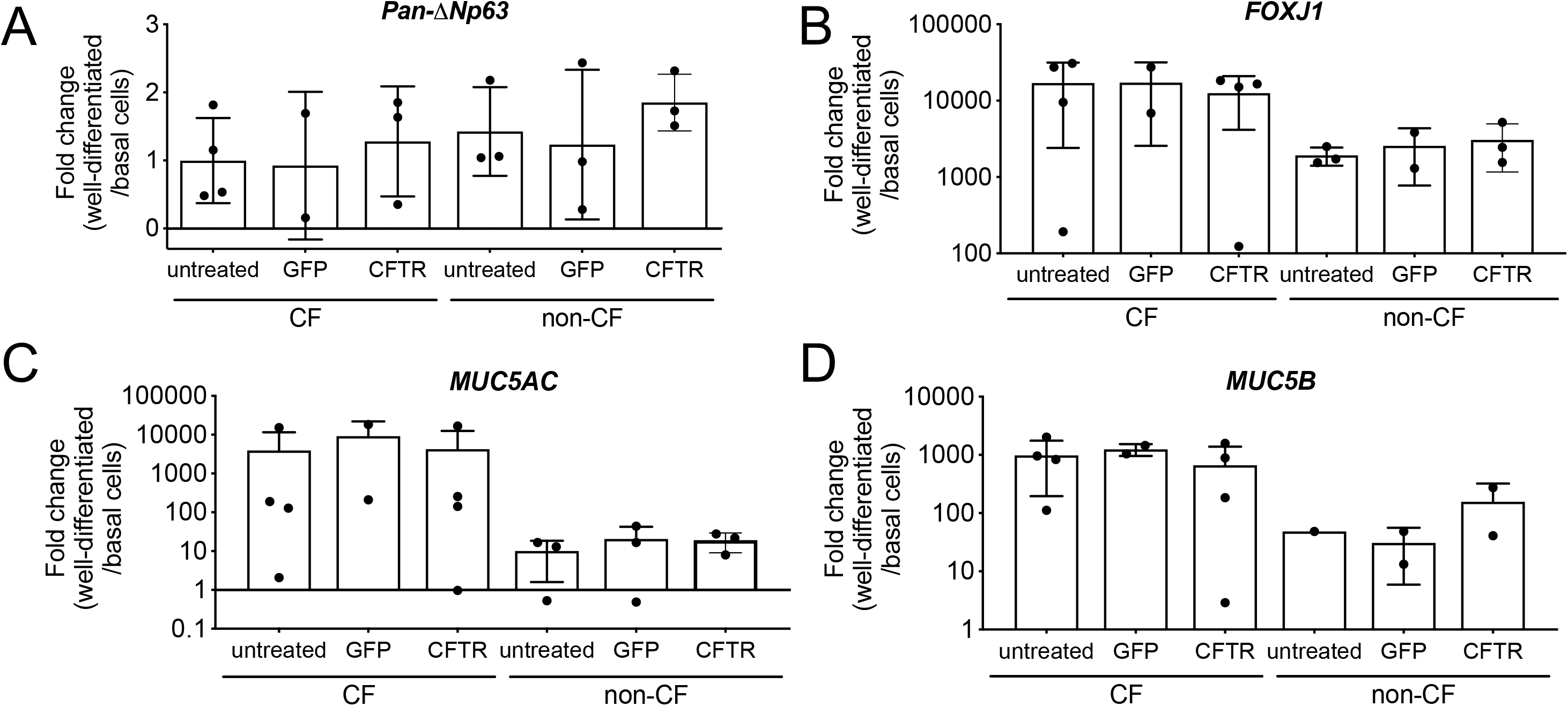
Transcript levels of known cell markers. Real Time quantitative PCR measurements were performed on basal cells and well-differentiated cultures. Basal cell marker *p63*, ciliated cell marker *FOXJI*, and secretory cell markers *MUC5AC* and *MUC5B* were quantified in all treatment groups. Fold change is expressed as well-differentiated over basal cells for each marker (n=5 CF donors, 4 non-CF donors).

**Supplemental Figure 3.**
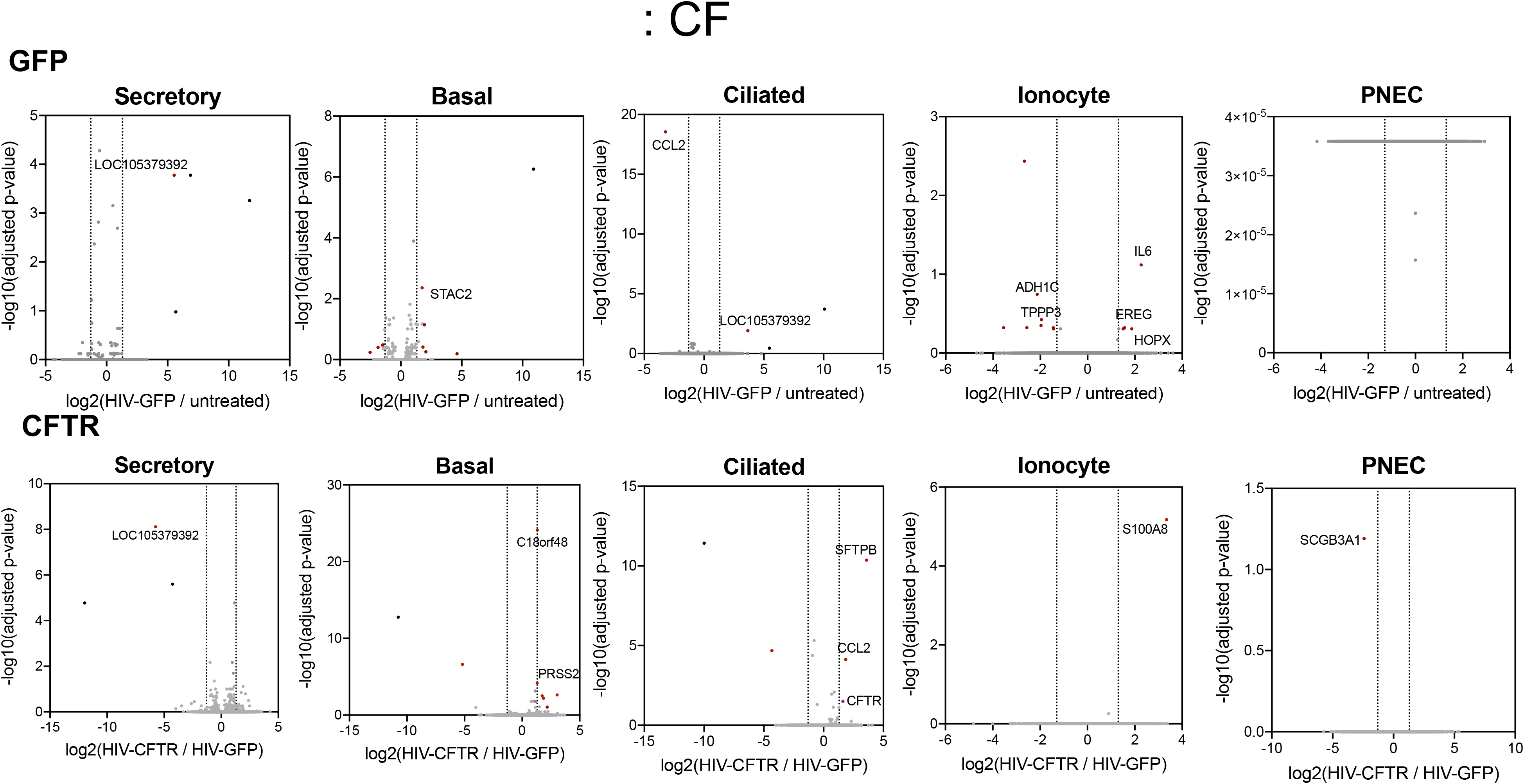
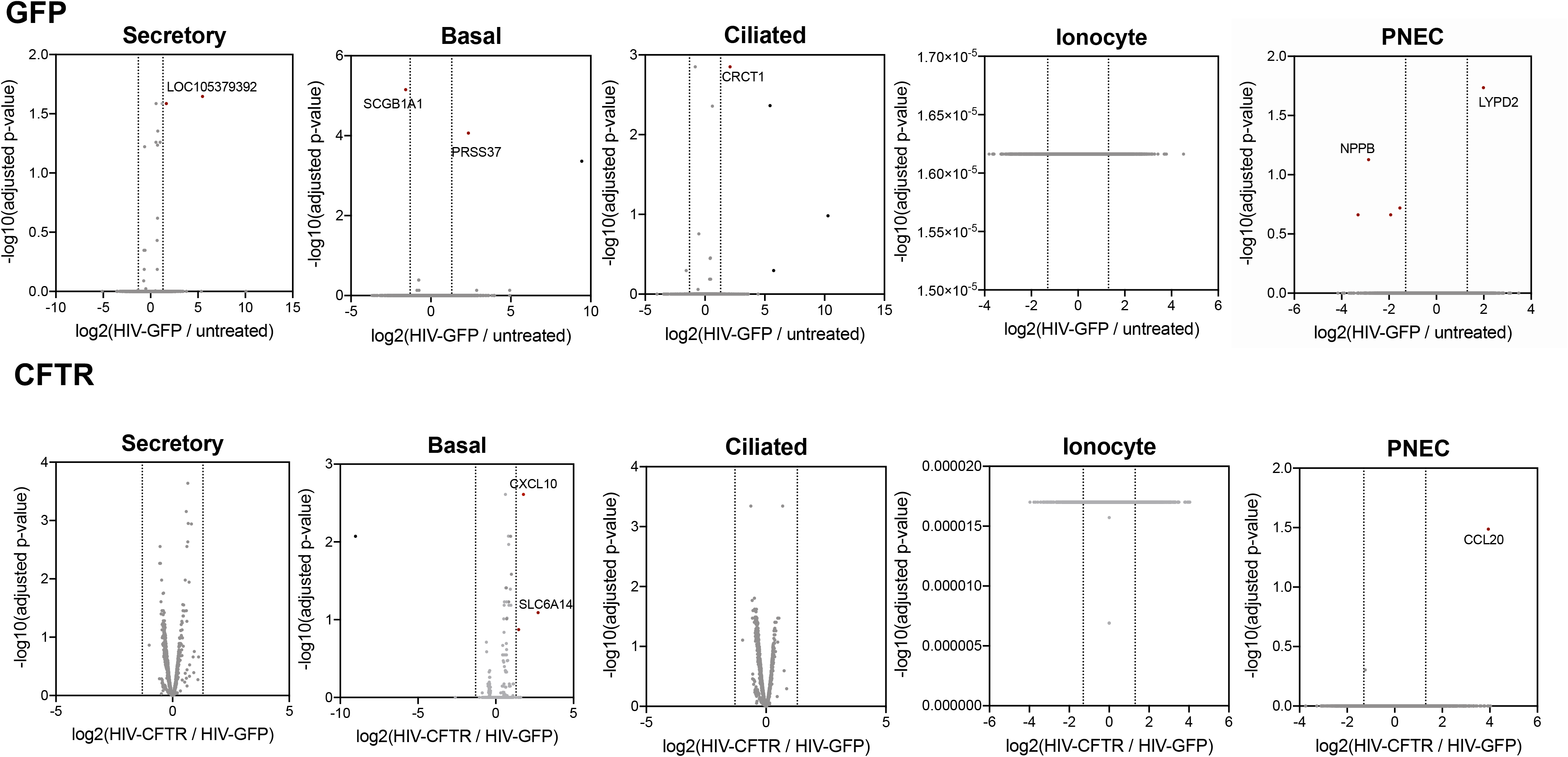
Effects of transduction and CFTR correction on cell-type specific gene expression. Differential expression analysis was performed to visualize the effect of HIV-GFP or HIV-CFTR transduction on gene expression for each cell type in CF and non-CF cells. Volcano plots show fold change in expression (log2 fold change) versus statistical significance (-log10(pval)). HIV-GFP gene expression (positive log2 fold change) was compared to untreated (negative log2 fold change). Red dots indicate differentially expressed genes, gray dots indicate non-differentially expressed genes, and black dots are vector encoded genes. For CFTR correction, gene expression was compared between HIV-CFTR (positive log2 fold change) and HIV-GFP (negative log2 fold change). Notable genes are labeled (n=4 CF, n=3 non-CF).

**Supplemental Table 1.** Genes with statistically significant changes in expression following exogenous CFTR delivery is shown. Genes increased or decreased after CFTR expression are represented as all cells combined as well as individual cell types (*FDR* <0.05).

|  | Cell type |  |  |  |  |  |
| --- | --- | --- | --- | --- | --- | --- |
|  | All | Basal | Ciliated | Secretory | Ionocyte | PNEC |
| ↑ after CFTR | CCL2, HES6, PRSS2, PLAU, IL32, KDR, CXCL14, SFRP1, KLK6, KLK8, LOC101929653, RASIP1, CLDN10 | C15orf48, CCL28, CCL2, PRSS2, PGBD5, IL23A, PLAU, IL32, PTN, KDR, IL34 | SFTPB, CCL2, PLAU, KRT17, CFTR | APOE, VIM, CCL2 | S100A8 |  |
| ↓ after CFTR | AZGP1, CYP4B1, DEGS2, SPIN4 |  | KRT4, AZGP1, FOLR1 | AZGP1, CYP4B1 |  |  |

**Supplemental Table 2.**
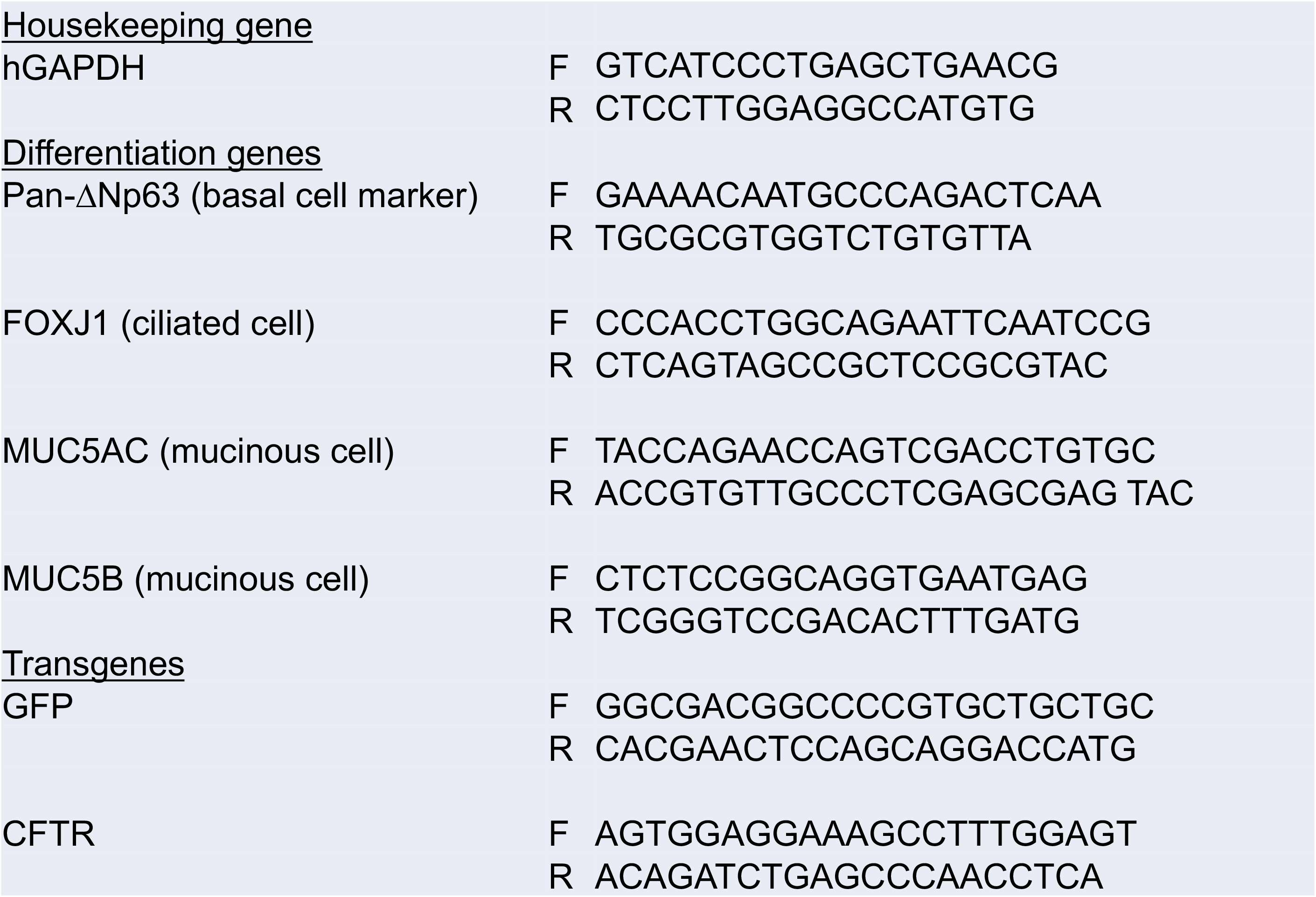
Primer sequences used in qRT-PCR assays are listed.

